# Genomic evidence for population-specific responses to coevolving parasites in a New Zealand freshwater snail

**DOI:** 10.1101/045674

**Authors:** Laura Bankers, Peter Fields, Kyle E. McElroy, Jeffrey L. Boore, John M. Logsdon, Maurine Neiman

**Author notes:** Corresponding author: Laura Bankers Department of Biology University of Iowa 143 Biology Building Iowa City, IA 52242, USA.

## Abstract

Reciprocal coevolving interactions between hosts and parasites are a primary source of strong selection that can promote rapid and often population- or genotype-specific evolutionary change. These host-parasite interactions are also a major source of disease. Despite their importance, very little is known about the genomic basis of coevolving host-parasite interactions in natural populations, especially in animals. Here, we use gene expression and sequence evolution approaches to take critical steps towards characterizing the genomic basis of interactions between the freshwater *snail Potamopyrgus antipodarum* and its coevolving sterilizing trematode parasite, *Microphallus* sp., a textbook example of natural coevolution. We found that *Microphallus*-infected *P. antipodarum* exhibit systematic downregulation of genes relative to uninfected *P. antipodarum*. The specific genes involved in parasite response differ markedly across lakes, consistent with a scenario where population-level coevolution is leading to population-specific host-parasite interactions and evolutionary trajectories. We also used an *F*_*ST*_-based approach to identify a set of loci that represent promising candidates for targets of parasite-mediated selection across lakes as well as within each lake population. These results constitute the first genomic evidence for population-specific responses to coevolving infection in the *P. antipodarum-Microphallus* interaction and provide new insights into the genomic basis of coevolutionary interactions in nature.

## Introduction

Host-parasite interactions constitute a primary source of natural selection and provide a powerful means of evaluating the evolutionary response to strong selection (Hamilton 1980; Abbate *et al.* 2015). Reciprocal antagonistic selection for greater host resistance and parasite infectivity can lead to antagonistic coevolution between host and parasite (Ehrlich & Raven 1964), which can in turn maintain high genetic diversity (Laine 2009; Bérénos *et al.* 2011; King *et al.* 2011; van Houte *et al.* 2016) and drive rapid evolutionary diversification (Laine 2009; Paterson *et al.* 2010; Masri *et al.* 2015).

While a great deal is known about the genomic underpinnings of plant-pathogen interactions (reviewed in Kover & Caicedo 2001), much less is known about animal systems, and particularly in natural populations (Routtu & Ebert 2015). Host-parasite interactions often have a genetic basis (*e.g.,* Paterson *et al.* 2010; Luijckx *et al.* 2013; Masri *et al.* 2015; van Houte *et al.* 2016), are linked to rapid molecular evolution (*e.g.*, Paterson *et al.* 2010; Tennessen *et al.* 2015), and are associated with gene expression changes (*e.g.,* Barribeau *et al.* 2014; de Bekker *et al.* 2015; McTaggart *et al.* 2015). Because most studies providing important insights into the genetic and genomic basis of host-parasite coevolutionary interactions have been limited to the laboratory (*e.g.,* Roger *et al.* 2008; Arican-Goktas *et al.* 2014; Barribeau *et al.* 2014; but also see Decaestecker *et al.* 2007; Routtu & Ebert 2015), we know very little about the evolutionary genomics of host-parasite interactions in natural populations (especially animals) and whether results from laboratory studies are recapitulated in nature (Routtu & Ebert 2015). As such, elucidating the processes of coevolution in nature is a requirement both to characterize a primary driver of selection and to formulate targeted strategies to fight natural parasite populations *(e.g.*, Gleichsner *et al.* 2015; Neafsey *et al.* 2015). Characterization of parasite-mediated changes in host gene expression provides a powerful means of deciphering the genomic basis of natural coevolutionary interactions because expression changes are often genotype or population-specific *(e.g.*, Barribeau *et al.* 2014), can affect host response to parasites (e.g., Brunner *et al.* 2013), and can underlie variation in resistance and susceptibility to infection (e.g., Stutz *et al.* 2015).

The New Zealand freshwater snail *Potamopyrgus antipodarum* and its sterilizing coevolving trematode parasite *Microphallus* sp. ("*Microphallus*") provide an especially compelling context for characterizing the genomic basis of coevolution in natural populations. First, selection imposed by both host and parasite is extremely strong: infected snails are completely sterilized (Winterbourn 1973), and *Microphallus* that cannot infect the first *P. antipodarum* by which they are ingested will die (King *et al.* 2011). Second, different *P. antipodarum* populations experience consistently high *vs.* consistently low *Microphallus* infection frequencies (Lively & Jokela 2002; King & Lively 2009; Vergara *et al.* 2013), indicating that infection frequency is a major determinant of the strength of parasite-mediated selection and, thus, the rate and mode of coevolution within each population. Third, *Microphallus* is locally adapted to *P. antipodarum* at both within- and among-lake scales (Lively *et al.* 2004; King *et al.* 2009), demonstrating the fine spatial scale of coevolution in this system. Fourth, the existence of many naturally replicated and separately evolving *P. antipodarum*-*Microphallus* interactions means that each population can be treated as a separate evolutionary experiment into the consequences of antagonistic coevolution (Dybdahl & Lively 1996). Finally, rapid coevolution of *P. antipodarum* and *Microphallus* has been documented in both a natural population (Jokela *et al.* 2009) and in an experimental coevolution study (Koskella & Lively 2009).

Multiple studies suggest that coevolutionary interactions between *P. antipodarum* and *Microphallus* fit a “matching alleles” infection genetics model, whereby there are no universally infective parasites or resistant hosts (Dybdahl & Krist 2004; Lively *et al.* 2004; Osnas & Lively 2005; King *et al.* 2009). Rather, *P. antipodarum* susceptibility depends on whether the genotype of the *Microphallus* individual matches the *P. antipodarum* individual at loci involved in resistance. Similar matching-allele mechanisms are thought to operate in other snail-trematode systems such as the laboratory model *Biomphalaria glabrata-Schistosoma mansoni* (reviewed in Adema & Loker 2015). Recent studies suggest that both allelic identity (Roger *et al.* 2008) and gene expression (Arican-Goktas 2014) are likely involved in mediating *B. glabrata* susceptibility to *S. mansoni* infection, emphasizing the potentially central role for gene expression in determining outcomes of host-parasite interactions in this and other snail-trematode systems (Hotez 2013; Jurberg & Brindley 2015).

Here, we take critical steps towards illuminating the genomic basis of coevolution in this textbook example of antagonistic coevolution (*e.g.*, Zimmer & Emlen 2013; Herron & Freeman 2014; Bergstrom & Dugatkin 2016) by using RNA-Seq to perform gene expression and *F_ST_* analyses for three replicated samples of *Microphallus*-infected *vs.* uninfected *P. antipodarum* from each of three different lake populations. In light of existing ecological evidence for population structure and local adaptation in this system, we expected to observe divergence between populations at both the gene expression and genetic level, such that population of origin as well as infection status would influence patterns of gene expression and relative levels of genetic differentiation. Such divergence at genetic and gene expression levels should be especially notable with respect to genes likely to be involved in the coevolutionary response to *Microphallus* infection.

## Materials and Methods

### Sample collection, dissection, and sample selection

Adult *P. antipodarum* were collected from shallow water (depth < 1m) habitats along the lake margins of three New Zealand lakes (Alexandrina, Kaniere, Selfe) known to contain relatively high *Microphallus* infection frequencies (~10-20%, Vergara *et al.* 2013; Table S1). Following transfer to the University of Iowa, we housed snails in 15 L tanks at 16°C with a 16:8 hour light:dark cycle and supplied *ad libitum* dried *Spirulina* (*e.g.*, Zachar & Neiman 2013) for one week to allow snails to recover from international travel prior to dissection and RNA isolation.

Because *P. antipodarum* is polymorphic for reproductive mode and ploidy level (Lively 1987; Neiman *et al.* 2011), we confined our RNA-Seq analyses to diploid adult non-brooding (non-reproductively active) females. These criteria ensured that we limited extraneous biological processes that may conflate differences in gene expression related to interactions between *P. antipodarum* and *Microphallus.* We dissected each snail to determine sex (male *vs.* female), *Microphallus* infection status (infected *vs.* uninfected), and reproductive status (brooding *vs.* non-brooding). We used the infection data to establish infection frequency for each lake sample (Table S1). Because *Microphallus* infection fills the body cavity of *P. antipodarum*, we confined analyses to head tissue. While the use of head tissue necessarily prevents the analysis of genes that are solely expressed in body tissue, this approach is the only way to ensure that comparable tissue types are isolated from both infected and uninfected snails.

All infected snails had stage five *Microphallus* infections (http://www.indiana.edu/~curtweb/trematodes/DATA_KEY.HTM). We first separated the head tissue of dissected snails, which does not contain *Microphallus* metacercariae, from body tissue, and then split the dissected head tissue into two halves. We immediately submerged one half of the head tissue in RNA*later*^^®^^ Solution (Life Technologies Corporation) and stored the tissue at 4°C for 24 hours followed by storage at −80°C (according to manufacturer protocol) until RNA extraction. The other head half was immediately snap-frozen in liquid nitrogen and stored at − 80°C for flow cytometric determination of ploidy level following Krist et al. (2014). For each RNA-sequencing replicate, we pooled head tissue from seven snails to obtain a sufficient amount of tissue for RNA extraction and sequencing. We obtained three biological replicates of parasite-infected and uninfected *P. antipodarum* from each of the three lakes, for a total of 18 replicates and 126 snails.

### RNA sequencing and *de novo* reference transcriptome assembly and annotation

We extracted RNA following the Invitrogen TRIzol protocol (Chomczynski 1993). RNA quantity and quality were assessed with a Bio-Rad Experion Automated Electrophoresis Station, following manufacturer protocol for the Experion RNA analysis kit with an RQI ≥ 8 and a minimum of 2µg of total RNA. RNA shearing and cDNA library preparation were completed following the Illumina Truseq LS protocol (Illumina, San Diego, CA, 2012). Following library preparation, we used an Illumina HiSeq 2000 for 2x100 bp paired-end RNA sequencing. Each RNA sample was given a unique indexed adapter sequence (Illumina, San Diego, CA, 2012) and was then separated into two halves, allowing us to sequence each sample twice in two different lanes in order to eliminate bias due to sequencing lane (two technical replicates per sample). We obtained a mean read length (SD) of 100.7 bp (0.96 bp) and a mean number of paired-end reads/replicated sample/lane (SD) of 10070420 (1474539.10), for ~20000000 paired-end reads/replicate. Next, we used FASTX Toolkit (Gordon & Hannon 2010) and FastQC (Andrews 2010) to trim adapters, assess sequencing quality, and filter out low-quality reads from the raw RNA-Seq data (see Table S2 for specific parameters). Quality filtering resulted in a mean sequence quality (Phred) score of 35.95, and a mean number of reads/replicate/lane (SD) of 9871440.4 (1468645.04), for a total of ~18000000 paired-end reads/replicate, about twice the coverage required to quantify expression differences across a wide range of expression levels (Wolf 2013).

We used Trinity v. 2.0.4 (Gabherr *et al.* 2011; Haas *et al.* 2013) to generate an initial *de novo* assembly using all of the combined filtered RNA-Seq data. First, we performed the recommended Trinity *in silico* normalization for each replicate in order to reduce the amount of memory required for the assembly process while maintaining a representative read set (following Gabherr *et al.* 2011; Haas *et al.* 2013). We then used Trinity to assemble the normalized reads (Haas *et al.* 2013), generating a transcriptome assembly with 462736 transcripts (Table S2). We annotated protein-coding regions in the transcriptome in order to identify long ORFs of putative genes that could be missed by homology searches and to filter out miscalled isoforms by using the Trinity plugin TransDecoder (Haas *et al.* 2013) (Table S2). This step also lessens the influence of sequencing errors, misassembled transcripts, chimeric sequences, and other common assembly issues (De Wit *et al.* 2015). Next, we used hierarchical clustering based on sequence identity to further reduce redundancy in the transcriptome assembly, as implemented in CD-HIT-EST (Huang *et al.* 2010) (Table S2). Finally, we identified and eliminated potential contaminant transcripts using the ‘blast_taxonomy_report.pl’ script from the Blobology pipeline (Kumar *et al.* 2013), blastx (Camacho *et al.* 2009) and a custom python script (available at: github.com/jsharbrough/grabContigs). We used Blobology to filter out transcripts with non-metazoan top blast hits, as well as Platyhelminthes (representing potential *Microphallus* contamination). These last filtering steps provided a final reference transcriptome assembly with 62862 transcripts.

We annotated the reference transcriptome by using blastx (Camacho *et al.* 2009) with an E-value cutoff of 1e-5. We then imported these blastx results into Blast2GO to assign gene ontology ("GO") terms to blastx-annotated transcripts (Conesa *et al.* 2005). We obtained blastx annotations for 10171 transcripts and both blastx and GO annotations for 15797 transcripts.

Nearly 75% of our transcriptome did not receive GO annotations, meaning that the functions of the majority of genes in our transcriptome cannot yet be determined. This result is unsurprising in light of a distinct deficit of mollusc genome sequence data available to aid in annotation (Kocot *et al.* 2015); though molluscs are the most species-rich animal phylum after arthropods (Smith *et al.* 2011), only ten (~1.4%) of the 700 sequenced animal genomes available on NCBI as of September 2016 are from molluscs.

### RNA-Seq gene expression analyses, functional enrichment, and *F_ST_* outlier analyses

We used our *de novo* transcriptome as our reference for all gene expression analyses. First, we used Tophat2 (Trapnell *et al.* 2013) to map filtered RNA-Seq reads to our *de novo* reference transcriptome. Next, we assembled mapped reads into transcripts and estimated transcript abundance with Cufflinks (Trapnell *et al.* 2012). We merged Cufflinks GTF and Tophat bam files for use in CuffDiff with Cuffmerge (Trapnell *et al.* 2012). Finally, we identified and quantified significant changes in gene expression using an FDR of 5% and Benjamini-Hochberg multiple test correction, as implemented in CuffDiff (Trapnell *et al.* 2012). We required that fragments per kilobase per million reads mapped (FPKM) exceeded zero to ensure that we were only comparing expression patterns among genes transcribed in all replicates (Table S2). We visualized the gene expression results with heatmaps, MDS plots, and dendrograms generated with cummeRbund (Goff *et al.* 2013), and used Blast2GO to compare the functions of differentially expressed genes and quantify the number of transcripts annotated with each GO term. We performed functional enrichment analyses and Fisher’s Exact Tests as implemented in Blast2GO to identify significantly overrepresented functional groups among differentially expressed genes (Table S2).

We used *F_ST_* outlier analyses to identify those genes with especially high levels of genetic differentiation between infected *vs.* uninfected snails; these genes provide a set of candidates for *Microphallus*-mediated selection. We filtered our reference transcriptome to include only transcripts with fragments per kilobase per million reads mapped (FPKM) > 0 in all 18 replicates (30685 transcripts) to ensure each replicate was represented for all loci in the *F_ST_* comparisons. We used Tophat2 (Trapnell *et al.* 2013) with the same parameters as in the gene expression analyses to map RNA-Seq reads to the filtered transcriptome. We then used Picard Tools to prepare mapped reads for variant discovery (http://picard.sourceforge.net), applying the AddOrReplaceReadGroups script to add read groups to each mapped bam file. We merged technical replicate bam files with the MergeSamFiles script, resulting in a single bam file for each of the 18 replicates. Finally, we used the MarkDuplicates script to identify duplicate reads, removing reads that mapped to more than one location in the transcriptome. Next, we used Samtools mpileup (Li *et al.* 2012) to call SNPs from processed bam files.

Analyses of levels of nucleotide variation from pooled RNA-sequencing data with programs like Popoolation2 (Kofler *et al.* 2011), which was developed to perform variant calling and calculate *F_ST_* from pooled sequencing data, can effectively identity SNPs and candidate focal genes of selection and perform *F_ST_-*based measures of genetic differentiation (*e.g.*, Fischer *et al.* 2013; Konczal *et al.* 2014). We used Popoolation2 (Kofler *et al.* 2011) to calculate *F_ST_* per site. The Popoolation2 pipeline begins by using the mpileup2sync.pl script to generate synchronized mpileup files that were filtered for base quality (Q20). We then applied the fst-sliding.pl script to calculate *F_ST_* per site. In all cases, we required a minimum coverage of ten, a maximum coverage of 200 (to reduce memory usage and biases introduced by inter-locus variation in gene expression), and that each SNP was sequenced at least four times (*e.g.*, Fischer *et al.* 2013; Konczal *et al.* 2014). We otherwise used default parameters. Finally, we filtered out SNPs with minor allele frequencies less than 5%. We chose these parameters to account for sequencing errors and to reduce the likelihood of obtaining false positives by ensuring that each SNP was sequenced multiple times, at the risk of not detecting rare alleles (*e.g.,* Konczal *et al.* 2014). We used IBM SPSS Statistics v. 23 to perform outlier analyses and identify outlier SNPs between infected and uninfected snails. We also identified the expression patterns of each transcript containing *F_ST_* outlier SNPs based on the expression analyses detailed above and evaluated whether transcripts containing *F_ST_* outliers between infected and uninfected snails contained outliers in multiple lake populations *vs.* only a single lake population. Finally, we compared relative levels of genetic differentiation (mean per-SNP *F_ST_* across all sites for which *F_ST_* could be estimated) between infected and uninfected snails within each lake and mean per-SNP *F_ST_* between lakes using Bonferroni-corrected Welch’s t-tests as implemented within IBM SPSS Statistics v. 23. The putative functions of transcripts representing *F_ST_* outliers were assessed using Blast2GO.

We used three complementary analytical approaches to evaluate the effect of infection and population on gene expression and genetic differentiation (Fig. S1). First, for the “inclusive analysis,” we identified differentially expressed genes likely to be broadly important for *P. antipodarum* response to *Microphallus* infection. We began by quantifying expression for each transcript for all pooled replicates under both conditions (infected and uninfected) to determine expression differences between all infected *vs.* all uninfected snails (Fig. S1a). We then compared the annotated functions of differentially expressed genes and performed functional enrichment analyses to determine the types of genes broadly important for infection response. Next, we calculated mean per-SNP *F_ST_* and performed *F_ST_* outlier analyses to measure relative levels of genetic differentiation within and between infected and uninfected snails and identified functional annotations and expression patterns of each transcript containing one or more *F_ST_* outlier SNPs.

Second, for the “within-lake analysis”, we compared gene expression patterns between infected *vs.* uninfected snails from each lake to characterize local (lake level) gene expression responses to *Microphallus* infection. This analysis included pairwise comparisons between infected and uninfected snails from within each lake population (Fig. S1b). We then compared the functions of differentially expressed genes between infected and uninfected snails within each lake. We also calculated mean per-SNP *F_ST_* and performed *F_ST_* outlier analyses between infected and uninfected snails within each lake. Finally, we determined the functional annotation and expression pattern of each outlier-containing transcript.

Third, in the “across-lake analysis”, we compared replicates by lake and infection status. Here, our goal was to use patterns of gene expression within and across lakes to evaluate evidence for local adaptation and identify genes likely to be locally important for response to *Microphallus* infection. We conducted every possible pairwise comparison of gene expression between all replicates from all three lakes (Fig. S1c), allowing us to differentiate between genes that were differentially expressed in infected *vs.* uninfected replicates across the three lakes (*i.e.*, evidence for population-specific infection responses) and genes expressed differently across lakes regardless of infection status (*i.e.*, expression difference due to lake of origin rather than infection status). Annotation and functional enrichment analyses allowed us to identify putative functions of genes that were significantly differentially expressed between infected and uninfected snails in more than one lake population *vs.* within a single lake. We visualized these comparisons with Euler diagrams generated with eulerAPE v3 (Micallef & Rodgers 2014). Finally, we evaluated whether transcripts containing *F_ST_* outliers between infected and uninfected snails contained outliers in multiple lake populations *vs.* only a single lake population.

## Results

### Inclusive analysis: Greater downregulation of transcripts in infected snails

We identified 1408 significantly differentially expressed transcripts (FDR: 5%, Benjamini-Hochberg) between *Microphallus-*infected and uninfected snails (Figs. 1a, S1a). Here, we define “upregulated” and “downregulated” as significantly higher and lower, respectively, expression in infected snails relative to uninfected counterparts. A significantly higher proportion of these 1408 transcripts were downregulated (1057, ~75%) *vs.* upregulated (351, ~25%) (Fisher’s Exact Test: *p* < 0.0001; Table 1, Fig. 1a), indicating that infected *P. antipodarum* experience systematic reduction in gene expression.

**Fig. 1.**
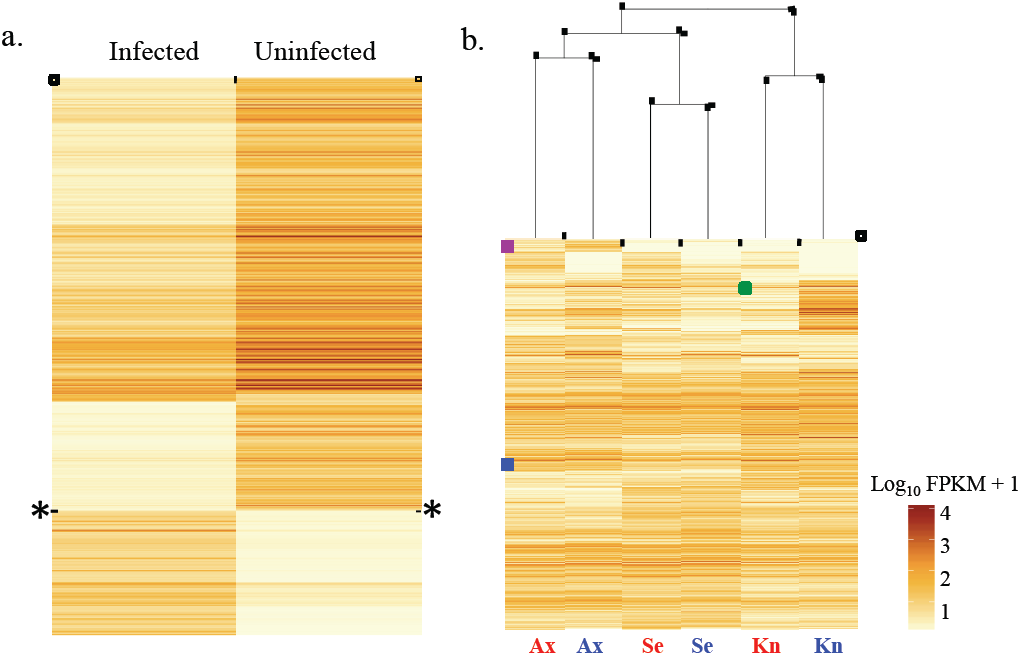
Heatmaps representing significantly differentially expressed genes. FPKM = fragments per kilobase per million reads mapped; higher FPKM = higher expression. Significance was assessed with a false-discovery rate of 5% and the Benjamini-Hochberg correction for multiple tests. a) Inclusive analysis: 1408 genes were significantly differentially expressed between infected and uninfected. Transcripts above the black line demarcated with “*” are significantly downregulated in infected snails (1057), and transcripts below this line are significantly upregulated in infected snails (351). b) Within-lake analysis: dendrogram and heatmap across infection status and lakes. “Kn Snails clustered by lake of origin rather than infection status, indicating spatially structured gene expression patterns.Grene box = an example of lake-specific expression regardless of infection status (downregulation in Alexandrina). Purple box = an example of genes that experience across-lake upregulation in infected snails (purple).

**Table 1.**
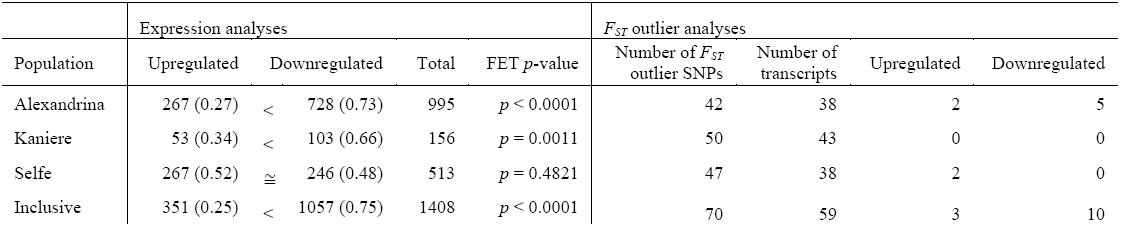
Summary of inclusive and within-lake analyses. Left: number of significantly upregulated, downregulated, and total number of significantly differentially expressed genes in infected *vs.* uninfected snails (proportion of differentially expressed genes). Right: number of *F_ST_* outlier SNPs between infected and uninfected snails, number of transcripts containing outlier SNPs, and the number of transcripts containing outlier SNPs that were significantly upregulated or downregulated in infected *vs.* uninfected snails.

We obtained gene ontology (GO) annotations for 447 of these 1408 transcripts, 216 of which were significantly upregulated and 231 significantly downregulated in infected snails. Sixteen of these upregulated genes were annotated as involved in immune system processes (Fig. S2), making these loci our strongest candidates for direct involvement in response to *Microphallus* infection. The ten genes with putative brain/behaviour functions that were upregulated in infected snails (Fig. S2) are particularly interesting candidates for future study in light of the well-characterized influence of *Microphallus* infection on *P. antipodarum* behaviour (*e.g.*, Levri 1999). Functional enrichment revealed that in infected snails, antigen processing and presentation were overrepresented among upregulated genes. Processes related to transcription and translation were overrepresented among downregulated genes, consistent with our observation that infected snails had significantly more downregulated genes (Table S3).

Our inclusive *F_ST_* analysis, comparing relative levels of genetic differentiation between infected and uninfected snails, revealed a mean per-SNP *F_ST_* (SD) of 0.0367 (0.0504) and identified 70 outlier SNPs (mean *F_ST_* of outliers (SD) of 0.3849 (0.2038)) from 59 transcripts. Of these 59 transcripts, ten were downregulated and three were upregulated in infected snails, meaning the majority of *F_ST_* outlier-containing genes were not significantly differentially expressed (Table 1). We obtained functional annotations for 16 of the 59 transcripts, four of which are likely involved in immune system processes (one transcript) and response to stimuli (three transcripts) (Tables S4, S5).

### Within-lake analysis: Lake-specific responses to *Microphallus* infection

Similar to the inclusive analysis, we observed a significantly greater proportion of downregulated *vs.* upregulated transcripts in infected *P. antipodarum* in two of the three lakes (Table 1, Figs. 1b, S1b). A total of 1539 transcripts were significantly differentially expressed between infected and uninfected snails in at least one of the three pairwise tests (Fig. 1b).

Functional enrichment for the within-lake analysis revealed that various metabolic processes are overrepresented among upregulated genes in infected snails from Alexandrina (Table S3). We did not detect significant functional enrichment for genes upregulated in infected snails from lakes Selfe or Kaniere. Upregulated genes in uninfected snails from Alexandrina and Selfe are enriched for GO terms related to transcription and translation. Upregulated genes in uninfected snails from Kaniere are enriched for pantothenate metabolism (Table S3).

The mean per-SNP *F_ST_* (SD) between infected and uninfected snails within each lake was 0.0485 (0.0504), 0.0479 (0.0623), and 0.0484 (0.0617) for Alexandrina, Kaniere, and Selfe, respectively (Fig. 2), providing evidence for population-specific levels of genetic differentiation between infected and uninfected snails. We identified 42, 50, and 47 *F_ST_* outlier SNPs between infected and uninfected snails (mean *F_ST_* of outliers (SD) of 0.3153 (0.1346), 0.3612 (0.1536), and 0.3352 (0.1442) from Alexandrina, Kaniere, and Selfe, respectively). Similar to the inclusive analysis, the vast majority of outlier-containing transcripts (93.5%) were not significantly differentially expressed (Table 1).

**Fig. 2.**
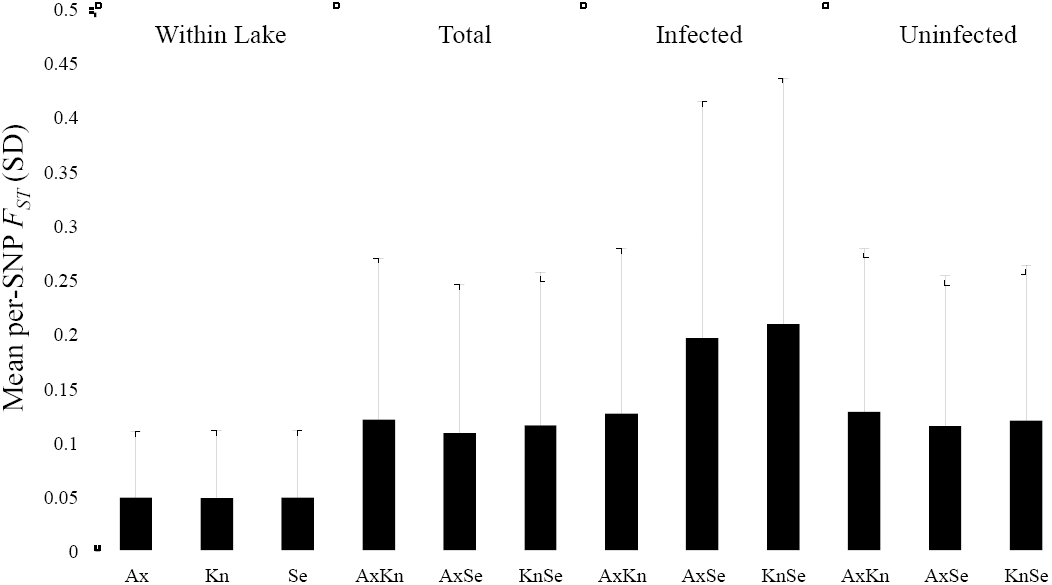
Mean per-SNP *F_ST_* (SD) calculated based on *F_ST_* per site. From left to right, the first panel is mean per-SNP *F_ST_* between infected and uninfected snails within each lake, the second panel is mean per-SNP *F_ST_* between lakes (*e.g.,* AxKn = mean per-SNP pairwise *F_ST_* between Alexandrina and Kaniere), the third panel is mean pairwise *F_ST_* for infected replicates only, and the fourth panel is mean *F_ST_* for uninfected replicates only. Lake acronyms follow Fig. 1. We used Welch’s t-tests to compare mean *F_ST_* (+/− SD) for all possible pairwise comparisons with Bonferroni multiple test correction (all within-lake infected *vs.* uninfected comparisons had significantly lower *F_ST_* than any of the across lake comparisons; *p* < 0.0001).

### Across-lake analysis: Lake of origin strongly influences gene expression and relative levels of genetic differentiation

We observed 6228 significantly differentially expressed transcripts within and across lakes (Figs. 3, S1c). ~75% (4689) of these 6228 transcripts were significantly differentially expressed across the three lakes regardless of infection status, indicating that lake of origin has a markedly stronger influence on gene expression than infection status (Figs. 1b, 3, 4, S3). Similarly, when replicates are clustered based on expression profile, they group first by lake of origin, followed by infection status. This result indicates both that population of origin is a key determinant of gene expression in *P. antipodarum* (Figs. 3, S3) and that infection results in predictable changes in expression (Fig. 3) that are still detectable on this background of population divergence. These findings are consistent with and extend to the gene expression level evidence for population-specific phenotypes (Holomuzki & Biggs 2006; Krist *et al.* 2014) and marked population genetic structure (Neiman & Lively 2004; Neiman *et al.* 2011; Paczesniak *et al.* 2013) in *P. antipodarum.* These results also demonstrate that both population of origin and infection have marked consistent and systematic consequences for gene expression in this species.

**Fig. 3.**
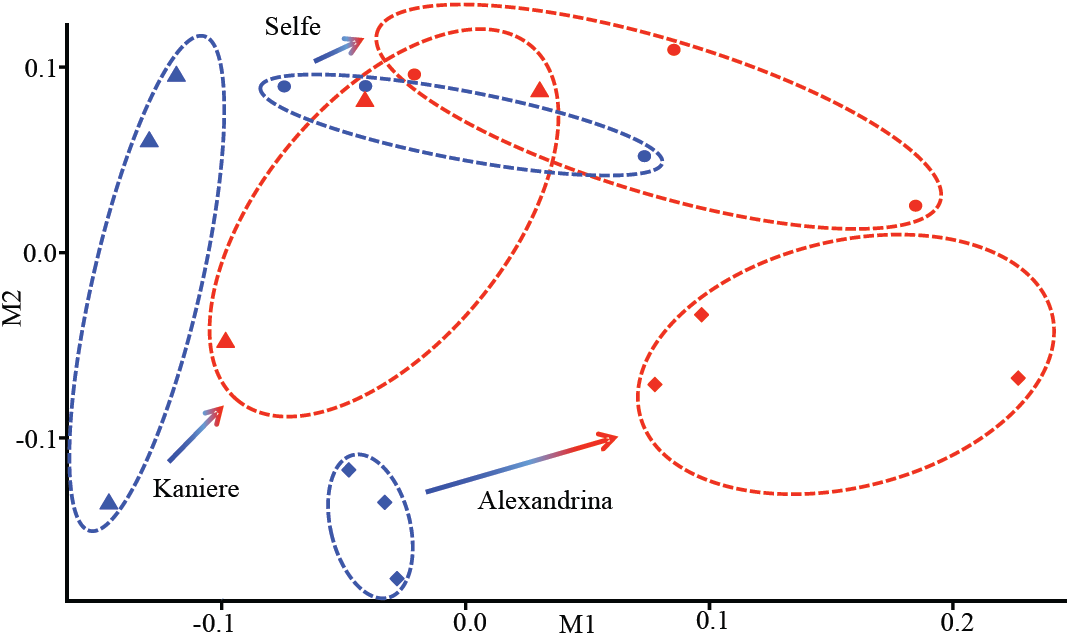
A multidimensional scaling (MDS) plot depicting how expression profiles are affected by lake population and infection status. Closer proximity of replicates in plot space indicates a more similar expression profile. Red = infected, blue = uninfected, diamonds = Alexandrina, triangles = Kaniere, circles = Selfe. The elliptical outlines (red = infected replicates from each lake; blue = uninfected replicates from each lake) delineate clustering for a particular replicate type and do not represent formal statistical tests; these clustering patterns are further supported by a dendrogram (Fig. S3). While the expression profiles that characterize uninfected and infected snails within each lake reveal lake-specific gene expression responses to infection, the consistent shift to the right along the “M1” axis for infected *vs.* uninfected snails for each lake population (highlighted with arrows) also demonstrates across-lake commonalities in expression profiles for infected snails.

We used the total number of transcripts identified as significantly differentially expressed between infected and uninfected snails (1539) in at least one of the three within-lake analyses (Fig. 1b) to compare the number of significantly up or downregulated transcripts in multiple *vs.* single populations, allowing us to determine whether snails from different lakes have similar gene expression responses to *Microphallus* infection. This comparison revealed that nearly all of the differentially expressed transcripts from the within-lake analysis (1447 transcripts, 94%) are only significantly differentially expressed in a single lake. Only 6% (92) of the differentially expressed transcripts showed significant differential expression in more than one population (Fig. 4). In summary, the vast majority of differentially expressed transcripts between infected and uninfected snails show significant up or downregulation in only one population, suggesting a distinct local (lake specific) gene expression response to parasite infection.

**Fig. 4.**
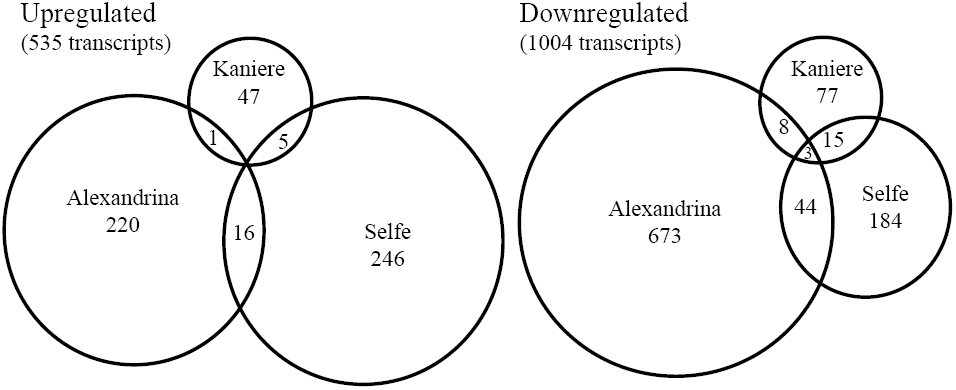
Euler diagrams depicting the number of significantly differentially expressed genes in infected *vs.* uninfected snails for the within-lake analysis. Left: number of upregulated genes in single *vs.* multiple lake populations (*e.g.*, 47 genes upregulated in Kaniere; 0 genes upregulated in all three lakes). Right: number of downregulated genes in single *vs.* multiple populations. Circle size is proportional to the number of genes in that category.

Next, we identified the types of genes that were significantly differentially expressed in single *vs.* multiple populations to parse out general *vs.* lake-specific infection responses. Of the 92 genes that were significantly differentially expressed in more than one lake, 22 genes were upregulated in infected snails (Table S6). We obtained GO annotations for seven of these genes, which had functions related to immune response, nervous system function, and metabolism. Ten of the 70 genes found to be significantly downregulated in infected snails in more than one lake received GO annotations, which included immune function, response to stimulus, and transcription/translation (Table S6). 396 of the 1447 genes that were significantly differentially expressed in only one lake received GO annotations (Fig. S2). Even though the particular genes experiencing differential expression differ on a lake-by-lake basis, these different genes often belong to similar GO categories (*e.g.*, immune system processes, response to stimulus, and behaviour).

We also evaluated how many transcripts contained *F_ST_* outlier SNPs in single *vs.* multiple populations. Similar to our gene expression analyses, we found that very few transcripts (six transcripts, ~5%) containing *F_ST_* outlier SNPs between infected and uninfected snails in one population also contained outlier SNPs in another population (three transcripts between Alexandrina and Kaniere, two transcripts between Alexandrina and Selfe, and one transcript between Kaniere and Selfe). This result indicates that the focal loci of *Microphallus*-mediated selection are likely to often be population specific. Of the transcripts that contained outlier SNPs, 12, 19, and 16 transcripts were annotated for Alexandrina, Kaniere, and Selfe, respectively. Two of these transcripts were annotated as relevant to immune system processes, 13 transcripts were annotated as involved in response to stimuli, and six transcripts were annotated with functions related to neurological/behavioural processes (Tables S4, S5).

Finally, we compared mean per-SNP *F_ST_* across lakes as a whole, for infected replicates only, and for uninfected replicates only. We found that *F_ST_* between infected and uninfected snails within each lake was significantly lower than the mean *F_ST_* for any across-lake comparison (Fig. 2), indicating relatively greater levels of genetic differentiation among lake populations than between infected and uninfected snails within a population. This result is consistent with and extends to the genome level the outcome of marker-based studies (*e.g.,* Paczesniak *et al.* 2013) that have documented strong across-lake genetic structure in *P. antipodarum.*

## Discussion

We used gene expression and *F_ST_* analyses to shed light on the genomic underpinnings of coevolution in a natural context. Most importantly, we found that population of origin is an important factor influencing gene expression, and that infection of *P. antipodarum* by its coevolving trematode parasite *Microphallus* elicits a marked, systematic, and population-specific gene expression response. These results have important implications for our understanding of the dynamics of coevolution in the *P. antipodarum*-*Microphallus* system, setting the stage for future research targeted at in-depth characterization of these population-level effects on the candidate loci that we identified. Our *F_ST_* analyses allowed us to evaluate relative levels of genetic differentiation within and among lakes and also revealed promising candidate genes for the focus of *Microphallus*-mediated selection. These findings provide a qualitative advance by extending evidence for local adaptation and coevolution in this textbook coevolutionary interaction to the genomic level, demonstrating distinct local genetic and gene expression responses by *P. antipodarum* to *Microphallus* infection. Together, these results illuminate the unique and important insights that can come from sampling multiple natural populations of interacting hosts and parasites.

### Systematic downregulation and response to parasite infection

Our inclusive analysis revealed that the majority of transcripts that are significantly differentially expressed in infected snails are downregulated relative to uninfected snails. This pattern could reflect several non-mutually exclusive phenomena, ranging from tissue/organ destruction and/or overall poor condition of infected snails (*e.g.,* Barribeau *et al.* 2014) to reallocation of resources to genes needed for defence against and response to *Microphallus* infection (*e.g.,* Ederli *et al.* 2015) to suppression and/or parasitic manipulation of *P. antipodarum* gene expression by *Microphallus* as a means of evading host immune and defence systems (*e.g.,* Levri & Lively 1996; Barribeau *et al.* 2014; Soper *et al.* 2014; de Bekker *et al.* 2015). Regardless of the specific mechanism(s) involved, the transcripts that are differentially expressed between infected and uninfected snails in the inclusive analysis represent a set of genes that are most likely to be of general importance to the *P. antipodarum*-*Microphallus* interaction. Our inclusive *F_ST_* analyses yielded additional candidate genes for the focus of parasite-mediated selection; the 70 SNPs from 59 genes that show relatively high levels of genetic differentiation provide a particularly strong set of candidate loci for *Microphallus* response. The presence of genes annotated with immune/stress responses and with neurological and behaviour-related functions strengthens this conclusion.

### Genomic evidence for population-level responses to infection

Our results demonstrate that *P. antipodarum* response to *Microphallus* infection is largely population specific. For example, in Alexandrina and Kaniere, the majority of significantly differentially expressed transcripts are downregulated in infected relative to uninfected snails (73% for Alexandrina and 66% for Kaniere). By contrast, the proportion of significantly upregulated and downregulated transcripts is much more similar in snails from Selfe (52% *vs.* 58%, respectively). This pattern is interesting in light of data from Vergara *et al.* (2013) suggesting that lake Selfe *P. antipodarum* have historically experienced lower frequencies of *Microphallus* infection (3.5x lower than Alexandrina, 1.4x lower than Kaniere; Table S1) (Vergara *et al.* 2013), suggesting that these populations may have experienced different historical intensities of *Microphallus*-mediated selection.

The vast majority (~87%) of differentially expressed transcripts were only significantly differentially expressed in a single lake population, and ~75% of significantly differentially expressed transcripts were expressed differently across lakes regardless of infection status, highlighting the important influence of population of origin on gene expression. We also found significantly lower levels of relative genetic differentiation between snails within *vs.* across lakes regardless of infection status, providing an intriguing potential connection between expression profile differences in different populations with genetic divergence between lakes. This result is consistent with previous evidence for population structure (*e.g.* Neiman *et al.* 2004; Neiman *et al.* 2011, Paczesniak *et al.* 2013) and/or negative frequency-dependent selection driving divergence among populations in the *P. antipodarum* system (*e.g.*, Dybdahl & Lively 1998). Together, these results demonstrate that lake of origin has a major influence on gene expression and relative genetic differentiation in *P. antipodarum* and provide the first genomic evidence consistent with lake-specific interactions between *Microphallus* and *P. antipodarum.*

### Candidate loci for future research

Our results provide a strong set of candidate loci for the genomic basis of *P. antipodarum* and *Microphallus* interactions. Three of the 16 upregulated immune-relevant genes are homologous to genes involved in response to trematode infection in the *Biomphalaria glabrata*-*Schistosoma mansoni* host-parasite system (*e.g.,* Mitta *et al.* 2005; reviewed in Coustau *et al.* 2015), suggesting these genes might play an important role in *P. antipodarum* response to *Microphallus* infection. We also identified seven downregulated genes with immune-related functions, including myosin light-chain kinase and fibrinogen-related proteins, which have also been shown to contribute to resistance to infection in the *Biomphalaria-Schistosoma* system (Mitta *et al.* 2005; Tennessen *et al.* 2015). Other significantly upregulated genes with immune-related functions in infected snails represent further candidates for response to *Microphallus* infection, while significantly downregulated genes with immune-related functions have the potential to be involved in resisting infection.

Among the genes upregulated in infected snails, we observed functional enrichment for actin, myosin, and genes with other cytoskeletal functions. Genes involved in cytoskeletal function, including actin and myosin, are upregulated upon parasite exposure in the *Biomphalaria*-*Schistosoma* system (reviewed in Coustau *et al.* 2015), indicating that these genes may also contribute to response to parasite infection in *P. antipodarum.* Oxidative processes have also been implicated in schistosome defence response in *Biomphalaria* (Adema *et al.* 2010, reviewed in Coustau *et al.* 2015); a preliminary line of evidence that oxidative processes may also be involved in the *P. antipodarum* trematode response is provided by significant functional enrichment of genes involved in oxidative processes in infected snails (*e.g.*, NADH oxidation, fatty acid beta-oxidation, energy derivation by oxidation of organic compounds).

The inclusive analysis revealed 21 upregulated and 16 downregulated genes with potential roles in important behavioural traits (*e.g.*, foraging, locomotion, mating) (Fig. S2). These results are consistent with evidence that exposure to (Soper *et al.* 2014) and infection by *Microphallus* (Levri & Lively 1996; Levri 1999) affects *P. antipodarum* behaviour. In particular, infected snails forage at a higher frequency than uninfected snails during the time of day when the waterfowl that are *Microphallus*’s final host are active, rendering infected snails more vulnerable to predation (Levri & Lively 1996). The implications are that these genes are a set of candidates for potential genetic mechanisms and pathways involved in *Microphallus*-induced alterations to *P. antipodarum* behaviour that could influence transmission probability. Future study of snails from natural populations featuring little to no *Microphallus* infection (Lively 1987; Lively & Jokela 2002) as well as manipulative experiments that allow comparisons between exposed *vs.* unexposed and infected *vs.* uninfected individuals will provide valuable additional steps forward by enabling differentiation between exposed but uninfected *vs.* naïve individuals.

Genes involved in the regulation of gene expression and ribosome structure and function were overrepresented among significantly downregulated genes. These results are consistent with our overall observation that infected snails had significantly more downregulated *vs.* upregulated genes, suggesting that *Microphallus* infection leads to decreased overall gene expression in *P. antipodarum.* These results are strikingly similar to a recent laboratory study showing that bumblebees (*Bombus terrestris*) exposed to particularly infective genotypes of a trypanosome parasite (*Crithidia bombi*) downregulated more genes than unexposed bumblebees (Barribeau *et al.* 2014). Similar results have been reported in other lab-based studies (*e.g.,* Roger *et al.* 2008; Arican-Goktas *et al.* 2014). Our results thus extend to natural coevolving populations the growing body of evidence that infected hosts experience systematic downregulation of gene expression.

### Summary and conclusions

We present novel evidence for systematic but population-specific genetic and gene expression responses to parasite infection in the *P. antipodarum*-*Microphallus* coevolutionary interaction. These results are the first genome-level evidence for the type of population-specific response expected under local adaptation and coevolution in this important host-parasite system. We also identified genes with functions related to immune and defence response and behaviour that are likely involved in the *P. antipodarum* response to *Microphallus* infection, providing a set of candidate genes for involvement in the genomic basis of coevolution. The targeted exploration of these genes made possible by these results will help illuminate the genetic and genomic mechanisms that determine the outcome of interactions between coevolving hosts and parasites in nature.

## Acknowledgements

We thank Gery Hehman for RNA sequencing assistance, Curt Lively and Daniela Vergara for field collections, Joel Sharbrough for the custom python script used in our transcriptome pipeline, and Kayla King, Thomas Städler, and Axios Review for helpful comments on an earlier version of this manuscript. The ploidy identification data presented herein were obtained at the Flow Cytometry Facility, which is a Carver College of Medicine/Holden Comprehensive Cancer Center core research facility at the University of Iowa. The Facility is funded through user fees and the generous financial support of the Carver College of Medicine, Holden Comprehensive Cancer Center, and Iowa City Veteran’s Administration Medical Center. This project was funded by NSF-MCB 1122176.

## Data Accessibility

Raw RNA sequencing reads will be deposited on the NCBI Sequence Read Archive. The annotated reference transcriptome assembly will be deposited on GenBank. Locations of sample sites, parameters for transcriptome assembly and analyses, and GO annotation and enrichment analyses are in the online supplementary information. The custom python script used in our transcriptome pipeline is available at: github.com/jsharbrough/grabContigs.

## Author Contributions

LB, MN designed research; LB performed research; LB, PF, KEM analysed data; LB, MN wrote the manuscript; JLB, JML, MN funded the study; LB, MN, PF, KEM, JLB, JML edited manuscript.

